# Engineered probiotics that sequester arsenite in a mouse gastrointestinal system

**DOI:** 10.64898/2026.01.16.699961

**Authors:** Nhu Nguyen, Miaomiao Wang, Sithembiso S. Msibi, Vincenzo Kennedy, Fateme Esmailie, L. Joseph Su, Lin Li, Clement T. Y. Chan

## Abstract

Chronic exposure to inorganic arsenic remains a major global health concern, as arsenite is frequently present in contaminated food and drinking water and readily absorbed through the gastrointestinal (GI) tract. Once internalized, arsenite accumulates in tissues and contributes to long-term health effects, including cancer, organ dysfunction, and neurological disorders. Despite extensive efforts to reduce environmental contamination, there are currently no practical strategies to prevent dietary arsenite from entering the human body during digestion. Here, we report a synthetic biology–based approach that uses engineered probiotics to detect and sequester arsenite directly within the GI tract before systemic absorption occurs. We engineered *Escherichia coli Nissle 1917* (*EcN*), a probiotic strain, to function as a living arsenite-interception system. Central to this design is an arsenite-responsive genetic toggle switch that activates chelator expression upon exposure and sustains production under biostatic conditions, while automatically shutting off during active cell division to limit metabolic burden and enhance biosafety. In parallel, we engineered an arsenite-binding protein derived from the transcriptional regulator ArsR to eliminate DNA-binding activity while retaining high-affinity metal binding, yielding a non-toxic chelator suitable for intracellular sequestration. The resulting engineered strain efficiently removed arsenite from its surrounding environment in vitro while maintaining robust cell viability and growth. To translate these findings to an in vivo context, we developed a mass-transfer model describing arsenite distribution among the stomach lumen, engineered bacteria, and epithelial cells. This model guided the selection of a bacterial dose predicted to substantially deplete lumenal arsenite prior to epithelial uptake. Using this strategy, we demonstrated in a mouse GI model that oral administration of engineered *EcN* markedly reduced arsenite entry into the bloodstream compared with wild-type EcN or no-bacteria controls. Together, these results establish a programmable probiotic platform for intercepting dietary arsenite and highlight a potential strategy for preventing absorption of environmental toxicants using living microbial therapeutics.

## Introduction

Heavy metal contamination represents a critical global health challenge. Inorganic arsenic, such as arsenite, is one of the most prevalent toxic metal species, with chronic exposure linked to cancers, organ failure, neurodegeneration, and other severe diseases (*1, 2*). As arsenite is hard to be eliminated once absorbed into human tissues, their effects are cumulative and long-lasting, and no safe threshold of exposure exists. Nonetheless, heavy metals remain abundant in human diets worldwide; more than 140 million people are estimated to consume arsenite-contaminated drinking water (*3*), millions in the United States are at risk from unsafe arsenite concentrations (*4*), and staple crops such as rice and wheat frequently contain high levels of arsenite (*5–7*). Despite the magnitude of this problem, there remains limited strategies to protect individuals from dietary exposures to this heavy metal species effectively. Thus, new approaches are urgently needed to mitigate this increasing health burden.

Current strategies to limit exposure focus on environmental management and monitoring. Approaches such as phytoremediation of contaminated soils (*8*), improved irrigation and waste practices (*9, 10*), and inspection programs supported by analytical techniques like mass spectrometry (*11, 12*) and atomic absorption spectroscopy (*13–15*) have value but are costly, infrastructure-dependent, and insufficient in regions with widespread contamination. Water purification systems can reduce exposure but remain inaccessible for many populations, and importantly, no practical technologies exist to prevent heavy metal uptake from food. Chelation therapy, while effective in acute poisoning, cannot be used preventively because of many adverse side effects caused by chelating agents (*16*). These limitations highlight an urgent need for affordable and scalable solutions to block dietary heavy metal absorption.

Microbial systems offer a promising foundation to tackle this health problem. Engineered bacteria expressing protein-based chelators such as metallothioneins (*17–19*), phytochelatins (*20*), and ArsR (*21*) can sequester toxic metals, and peptides with high binding affinity have been designed for similar purposes (*22–24*). In parallel, transcriptional regulators that sense specific metals have been harnessed for biosensing applications (*25–28*); recent innovations in directed evolution (*29, 30*) and genetic circuit amplification (*31*) have shown promising paths to expand sensitivity. However, most efforts have centered on detection rather than incorporating genetic sensing with biological chelators to promote heavy metal removal.

Emerging synthetic biology approaches have been explored to engineer gut microbes as versatile living therapeutics. Probiotic strains have been developed to metabolize harmful molecules, suppress pathogens, and secrete bioactive molecules to improve host physiology (*32–37*). These advances demonstrate that gut microbes can be designed to sense environmental or host-derived signals and deliver therapeutic responses in situ.

Here, we have explored the concept of microbial sense-and-response for an environmental toxicology application by engineering gut bacteria to detect and sequester arsenite in the gastrointestinal tract before absorption occurs. A key advantage of engineered bacteria is their ability to couple sensing and intervention in real time. Unlike conventional chelator therapies, which bind both toxic and essential metals, engineered microbes can be designed to activate metal-binding only when a target contaminant is present, thereby reducing off-target effects and minimizing disruption of essential metal homeostasis. This dynamic, conditional response also allows bacteria to conserve resources for their cell fitness in the drastically changing gastrointestinal environment yet rapidly increase sequestration capacity during exposure events. Together, these features provide a unique platform for selective, preventative detoxification that is not achievable with existing chemical or physical methods.

Specifically, we have developed an *Escherichia coli Nissle 1917* (*EcN*) strain equipped with an arsenite-responsive genetic circuit to control an output module to express a protein-based arsenite chelator. We demonstrated that this engineered bacterial strain performed in a mouse gastrointestinal system to reduce the rise in blood arsenite levels after an oral administration of arsenite solution. This work establishes a new preventative strategy against dietary metal exposure and broadens potential applications of engineered microbes to include protection from environmental toxicants.

## Results

### Design of genetic circuit for sensing and responding to arsenite

For protection against dietary arsenite, we designed an engineered strain of *EcN* that can pass through the digestive tract, capture the toxic metal, and be excreted while retaining the bound arsenite. The strain was designed to express a protein-based arsenite chelator, enabling bioaccumulation of arsenite-protein complexes within bacterial cells. A key feature of our design is that the bacteria will produce a basal level of chelator even in the absence of heavy metals, ensuring they are primed to capture toxicants as soon as exposure occurs. This basal level is intentionally kept low to minimize non-specific binding of essential metal ions and to preserve cellular fitness. Upon encountering arsenite, the genetic circuit triggers a robust induction of chelator expression, dynamically reallocating cellular resources to maximize sequestration at the point of highest toxicant burden. Notably, the circuit is further designed to sustain elevated chelator expression in arsenite-exposed cells even after arsenite levels decline, ensuring effective clearance of residual toxicants and extending the protective effect during digestion.

The engineered *EcN* cells are intended for repeated oral administration and most cells will be eliminated from the gastrointestinal tract after passage; however, a small fraction may persist and colonize the gut for extended periods (*38*). To prevent these proliferating cells from uncontrollably producing the chelator in the absence of arsenite, we incorporated a growth-dependent control feature. When arsenite concentrations fall below the detection threshold, chelator synthesis continues only in non-dividing cells; active cell division over several cycles leads to the cessation of chelator expression.

This event-triggered, sustained response is enabled by a genetic toggle switch architecture (**Figure 1**). Although the ArsR transcriptional regulator functions as an arsenite sensor (*39*), directly coupling output expression to ArsR activity restricts both sensitivity and the duration of response. To overcome these limitations, we implemented reciprocal repression between ArsR and a counter-regulator, CelR—a transcriptional repressor that controls gene expression from an engineered promoter (*40*). In the absence of arsenite, ArsR dominates the regulatory circuit and represses both the chelator and CelR. Upon arsenite exposure, ArsR repression is relieved, allowing expression of both genes; newly synthesized CelR then suppresses ArsR, reinforcing the ON state. This architecture confers two key advantages. First, it enables a prolonged response: because CelR continues to inhibit ArsR after arsenite levels fall, chelator expression persists, supporting clearance of residual toxicants. Second, it enhances sensitivity: by lowering ArsR abundance through CelR-mediated repression, the circuit reduces the arsenite threshold required to fully switch into the ON state.

**Figure 1.**
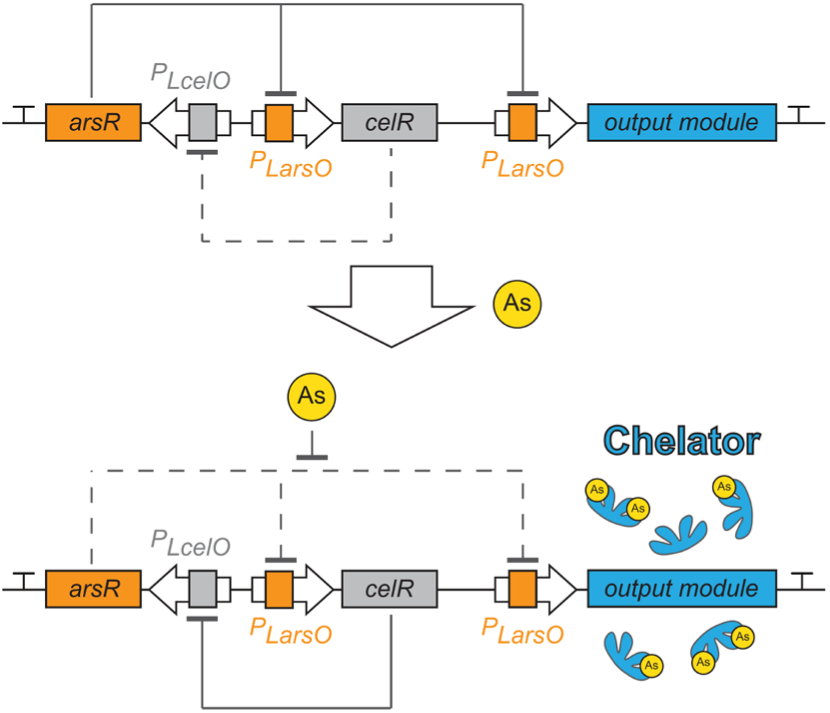
Circuit design for sensing and responding to arsenite. The circuit consists of two regulatory proteins, ArsR and CelR, that form a toggle switch. Under normal growth conditions without arsenite, ArsR is highly expressed and dominant, maintaining the circuit in the chelator- OFF state. Upon arsenite exposure, CelR and chelator expressions are induced, which represses arsR transcription. Once triggered, chelator expression continues even after arsenite is removed, until active cell division reduces CelR levels and restores ArsR expression, returning the circuit to the OFF state.

Importantly, the circuit also incorporates a mechanism that ties output decay to active cell division. We intentionally limited the maximal intracellular CelR level so that rapid growth dilutes CelR faster than it can repress ArsR, allowing ArsR to regain dominance and gradually return the circuit to the OFF state. Together, these design features transform a transient environmental cue into a durable yet self-limiting detoxification program, establishing a programmable strategy for microbial heavy-metal sequestration in vivo.

### Arsenite-responsive toggle switch circuit

To establish the design in **Figure 1** for desired arsenite sensing and response behaviors, we first constructed the Control circuit shown in **Figure 2A** to characterize the transcriptional regulator ArsR as an arsenite-responsive genetic sensor; the circuit was built in a plasmid that can be hosted by *EcN*. The *arsR* gene was placed under the control of the promoter *P_LcelO_* (*40*); transcriptional activity of this promoter is repressed by the regulator CelR. In the absence of CelR, ArsR was therefore constitutively expressed in this circuit. ArsR specifically binds to the *arsO* operator (*39*); we incorporated this operator downstream of the *P_L_* promoter (*41*) to generate *P_LarsO_* (**Supplementary Figure S1**). Binding of ArsR to this operator represses transcription from the engineered promoter. We next characterized the signal-response behavior of the *arsR*/*P_LarsO_* system using mCherry as a fluorescent reporter. *EcN* cells harboring this circuit were exposed to a range of sodium arsenite concentrations (0 to 5 µM) overnight, and mCherry fluorescence was quantified by flow cytometry (**Figure 2B**). In unexposed cells, mCherry fluorescence averaged 975 ± 41 a.u. (mean ± S.D.), whereas exposure to 5 µM arsenite increased fluorescence to 12,237 ± 1,511 a.u., confirming that the system is responsive to arsenite.

**Figure 2.**
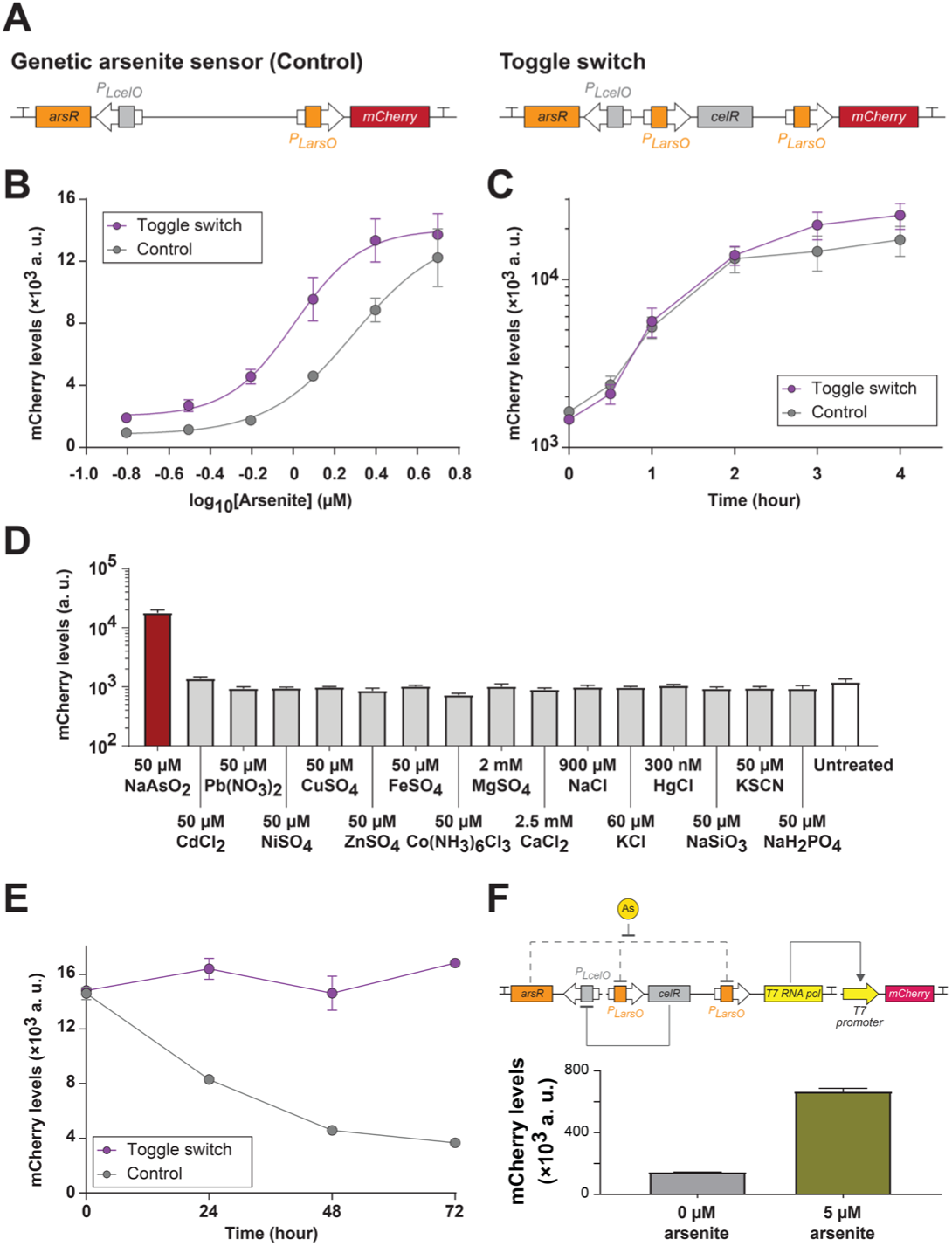
Characterization of the arsenite-responsive toggle switch circuit. (**A**) The circuit design of the Control and Toggle Switch circuits are shown. (**B**) Sensitivity of these two circuits were assessed by measuring cellular response to a 24-hour exposure to 0 to 5 μM arsenite. The equation of illustrated dose-response curves is presented in the Methods section. (**C**) To evaluate the rate of output production, cellular levels of mCherry were measured in 0 to 4 hours of exposing to 5 μM of arsenite. (**D**) Specificity of signal detection was assessed by exposing cells containing the toggle switch circuit to various conditions with different chemical species and measuring changes in cellular mCherry levels. (**E**) Stability of output production was characterized; arsenite-induced cells were transferred to a biostatic condition without arsenite for 72 hours with measurement of cellular mCherry levels every 24 hours. (**F**) To enhance output production levels, the characterized Toggle Switch circuit was used to control the expression of T7 RNA polymerase for driving mCherry production as the output. *EcN* with this circuit was incubated with 0 and 5 μM of sodium arsenite overnight before characterizing cellular mCherry levels. Each data point represents mean ± S.D. of three biological replicates.

Next, we constructed the toggle switch by incorporating the *celR* regulator gene into the plasmid, placing it under the control of the *P_LarsO_* promoter so that ArsR and CelR mutually repressed each other’s expression (Toggle Switch circuit; **Figure 2A**). Our goal was to develop a sense-and-response system in which exposure to arsenite triggers expression of the output gene, and once activated, the output remains expressed even after arsenite is removed, until cells resume active division during growth. Similar growth-dependent toggle switch behavior has been developed previously (*42*).

To achieve this functionality in our arsenite-sensing circuit, we fine-tuned CelR expression by modulating its ribosome binding site (RBS) strength across a range of predicted translation efficiencies (*43*). Each circuit variant was evaluated to identify the version that produced the desired response dynamics (**Supplementary Figure S1**). For initial screening, cultures were grown overnight in the presence of 0 or 5 µM sodium arsenite to allow full induction of output expression. We sought a circuit variant that exhibited low mCherry expression in the absence of arsenite and strong mCherry induction upon arsenite exposure, consistent with the expected toggle switch behavior. To identify circuit variants capable of sustaining output expression under biostatic conditions after arsenite removal, cells from 5 µM arsenite-exposed cultures were collected and resuspended in an equal volume of fresh culture medium lacking arsenite, thereby maintaining cell density and ensuring a stationary-phase state. These cultures were then incubated under standard growth conditions for 24 hours before mCherry fluorescence was measured.

The mCherry expression levels varied across cells harboring different circuit variants (**Supplementary Figure S1**). When the RBS driving *celR* was too strong (e.g., RBS 1), mCherry fluorescence was high under both 0 and 5 µM arsenite conditions, indicating that excessive CelR expression suppressed *arsR* transcription regardless of arsenite presence. Among the variants exhibiting differential responses between 0 and 5 µM arsenite (RBS 2 to 13), most failed to maintain mCherry expression after arsenite removal for 24 hours. In the Control circuit lacking *celR*, mCherry fluorescence decreased from 14,600 ± 367 to 8,297 ± 105 a.u. after this biostatic incubation; several toggle switch variants displayed similar behavior, suggesting that these circuits produced insufficient CelR levels to repress ArsR even after arsenite induction. In contrast, the circuit containing RBS 10 maintained high mCherry fluorescence under biostatic conditions following arsenite removal. This variant satisfied our design criteria and was therefore selected for subsequent characterization.

### Characterization of circuit response

We compared the arsenite sensitivity of this selected toggle switch circuit with that of the control circuit (**Figure 2B**). Cultures harboring either construct were exposed overnight to a range of sodium arsenite concentrations, and cellular mCherry fluorescence was quantified. The resulting data were fitted to dose-response curves, revealing that the toggle switch enhanced sensitivity to arsenite relative to the control. Specifically, cells carrying the toggle switch reached half-maximal mCherry levels (EC_50_) at 1.02 µM arsenite, whereas those with the control circuit required 1.97 µM. Furthermore, the Hill slope of the toggle switch response (2.71) was steeper than that of the control circuit (2.09), indicating a more pronounced, cooperative increase in output expression with rising arsenite concentrations. This behavior is consistent with the expected mechanism in which arsenite-induced CelR expression suppresses ArsR, thereby amplifying mCherry expression.

Additionally, we monitored how quickly the circuits responded to arsenite exposure (**Figure 2C**). Both engineered strains were treated with 5 µM sodium arsenite at the mid-log phase, and their mCherry fluorescence was measured over time. A clear increase in fluorescence was detected within the first hour in both cases, indicating that the regulator ArsR rapidly sensed and responded to arsenite. After the initial response, mCherry levels in both strains continued to rise gradually.

We next evaluated the specificity of the toggle switch for arsenite detection (**Figure 2D**). In the gastrointestinal environment, the engineered bacteria are likely to encounter a wide variety of ionic species; therefore, it is essential that the circuit responds selectively to arsenite to avoid unnecessary chelator production, which could waste cellular resources and lead to off-target metal sequestration. Engineered *EcN* carrying the toggle switch was exposed to different chemicals containing various cations and anions. Among all tested conditions, cells exposed to 50 µM sodium arsenite exhibited a marked increase in mCherry fluorescence (approximately 15-fold), whereas cells treated with other ions showed changes of less than 2-fold. These results demonstrate that the toggle switch responds specifically to arsenite and is largely insensitive to other physiologically relevant ions.

Furthermore, we examined the stability of output expression after arsenite removal in the engineered *EcN* culture (**Figure 2E**). To prevent uncontrolled chelator production during active growth, our design aimed for a circuit that sustains output expression only under biostatic conditions and halts production when cells resume division. To maintain arsenite-induced cells in a non-growing state, saturated cultures were collected daily and resuspended in an equal volume of fresh medium, replenishing nutrients for cell viability without dilution. This approach preserved high cell density and maintained cells in a biostatic phase. Under these conditions, cells carrying the Control circuit exhibited a sharp decline in mCherry fluorescence after arsenite removal, whereas those harboring the Toggle Switch maintained elevated fluorescence for at least three days. This sustained expression is consistent with CelR-mediated maintenance of the ON state, indicating that the toggle switch confers a form of molecular memory that prolongs output production in the absence of continued arsenite exposure.

In contrast, resumption of cell growth triggered the toggle switch to revert to the OFF state (**Supplementary Figure S2**). To examine the effect of cell division on circuit stability, we diluted the cultures 2-, 3-, or 6-fold after each medium exchange, thereby modulating the number of division cycles required for the cultures to reach saturation. In toggle switch cultures subjected to 2-fold daily dilution, mCherry fluorescence decreased only about 2-fold after three days, indicating that output production persisted but at a reduced rate. However, higher daily dilution ratios (3- and 6-fold) led to a more pronounced decline in mCherry levels, consistent with active cell division resetting the circuit to the OFF state. This behavior highlights a key safety feature of the design—ensuring that proliferating cells in the host do not continuously produce excessive levels of the chelator.

To further enhance output production, we coupled the Toggle Switch circuit to a T7 RNA polymerase-driven expression module, enabling transcriptional amplification of the output gene (**Figure 2F**). The translation efficiency of T7 RNA polymerase was optimized by tuning its RBS strength (**Supplementary Figure S3**), with sequences designed using the computational RBS calculator (*43*). Each circuit variant was induced with arsenite for 1 hour, and cellular mCherry fluorescence was quantified. Among the tested variants, the construct containing RBS C exhibited the greatest dynamic range, with mCherry fluorescence increasing from 54,481 ± 1,409 a.u. (uninduced) to 235,523 ± 7,685 a.u. (induced). When cultures of this variant were incubated overnight to assess maximal output, fluorescence reached 143,220 ± 1,489 a.u. and 664,975 ± 18,216 a.u. in uninduced and induced cells, respectively. These results demonstrate that coupling the Toggle Switch to a T7-based transcriptional amplifier markedly boosts output expression. The strong, orthogonal activity of T7 RNA polymerase likely insulates the output expression from host transcriptional system, thereby expanding its dynamic range and enhancing signal fidelity.

### Development of an arsenite chelator

ArsR is a transcriptional regulator that responds to arsenite as an inducer and belongs to the ArsR/SmtB family (**Figure 3A**). An ArsR homolog binds arsenite with a dissociation constant of approximately 10^-15^ M, indicating exceptionally tight binding (*44*). Based on this property, we selected ArsR as the chelator for our engineered bacteria. However, high-level expression of ArsR proved toxic to *E. coli* as we demonstrated that overexpression of this gene for three hours under an inducible promoter caused roughly a 2-log reduction in colony-forming units (CFU; **Figure 3B**). Consistent with this observation, previous studies have also reported growth inhibition upon ArsR expression (*45*). We hypothesize that this cytotoxicity arises from non-specific binding of ArsR to genomic DNA; at high intracellular concentrations, ArsR may repress essential genes, thereby compromising cell viability.

**Figure 3.**
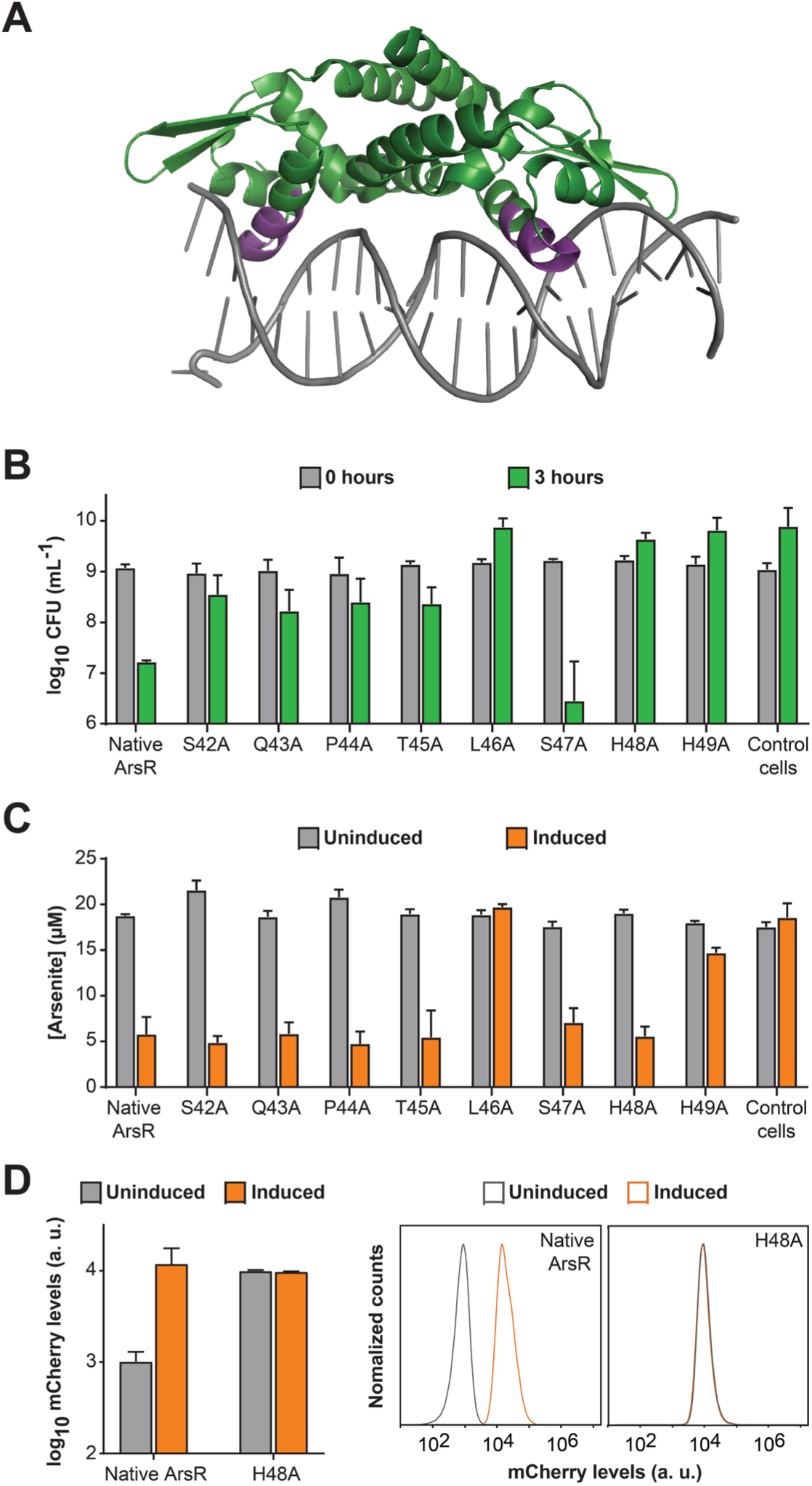
Development of the arsenite chelator. (**A**) Structure of NolR, a ArsR/SmtB family regulator, in complex to a DNA oligonucleotide (PDB:4OMY). The homologous region in α4 (residues 42 to 49) for mutation is colored purple, which situated in the DNA major groove. (**B**) Assessment of cytotoxicity of ArsR mutant expression to *E. coli*. Colony-forming units in each culture was measured before induction (0 hours) and three hours after induction (3 hours). (**C**) Efficiency of arsenite sequestration. Cultures with and without induction for the expression of an *arsR* mutant were incubated with 20 µM sodium arsenite for 3 hours before collection of supernatants for arsenite quantification. (**D**) Characterization of ArsR derivatives in controlling gene expression. ArsR and ArsR H48A were used to control mCherry expression in *E. coli*. After overnight induction with 5 µM arsenite, cells were collected to measure mCherry levels with flow cytometry (left panel). Representative raw data were presented on the right. Each data point represents mean ± S.D. of three biological replicates.

To test this hypothesis and to develop an arsenite chelator with reduced cytotoxicity, we sought to engineer an ArsR mutant that retains arsenite-binding ability but loses DNA-binding capacity. Members of the ArsR/SmtB family contain a winged helix-turn-helix DNA-binding motif, with helices α3 and α4 directly contacting the major groove of DNA (*46, 47*). The N-terminal region of α3 is also involved in metal chelation (*48, 49*). Our strategy was to disrupt DNA binding while minimizing perturbation of the metal-binding site. To this end, we introduced single amino acid substitutions within residues 42 to 49 of helix α4 of ArsR, a region predicted to mediate DNA interaction but not metal binding based on the protein-DNA complex structure of a family member, NolR (**Figure 3A**), and sequence alignment among ArsR/SmtB family regulators (**Supplementary Figure S4**).

Among the ArsR mutants generated, individual amino acid residues were substituted with alanine. Each *arsR* mutant gene was expressed under the control of a T7 promoter in an *E. coli* strain harboring an arabinose-inducible T7 RNA polymerase gene in its genome. To evaluate cytotoxicity, we compared the colony-forming units (CFUs) of cultures with and without induction of *arsR* expression by 0.2% arabinose. Notably, most substitutions alleviated cytotoxicity, except for S47A (**Figure 3B**). Three mutants, including L46A, H48A, and H49A, restored cell growth to levels comparable to the control strain lacking ArsR expression, as determined by CFU measurements.

In parallel, we assessed the arsenite sequestration capacity of these strains. Induced and uninduced cultures with each mutant were incubated with 20 µM of sodium arsenite, after which supernatants were collected to quantify residual arsenite levels (**Figure 3C**). Among the three mutants with the lowest toxicity, L46A and H49A mutants exhibited reduced arsenite removal efficiency, whereas the H48A mutant maintained strong arsenite-binding capacity, achieving arsenite reduction comparable to that of native ArsR. Therefore, H48A was selected as the optimized arsenite chelator for subsequent studies.

To confirm the loss of DNA-binding function in the H48A mutant, we evaluated its ability to regulate an ArsR-responsive promoter controlling mCherry expression in *E. coli* (**Figure 3D**). The genetic design, shown as the Control circuit in **Figure 2A**, includes a constitutively expressed *arsR* gene that represses mCherry expression as a reporter. In cells expressing native ArsR, mCherry fluorescence increased approximately 10-fold upon arsenite exposure. In contrast, when the native *arsR* gene was replaced by the *arsR H48A* mutant, cells expressing the ArsR H48A mutant exhibited high mCherry expression even in the absence of arsenite, and fluorescence levels remained unchanged after induction. These results indicate that ArsR H48A has lost its DNA-binding capability, consistent with its reduced cytotoxicity.

### Engineered *EcN* for arsenite sequestration

After establishing the genetic sense-and-response circuit (**Figure 2**) and the arsenite chelator module (**Figure 3**), we combined these components to construct the final engineered *E. coli Nissle 1917* (*EcN*) strain capable of detecting and sequestering arsenite. In this design (**Figure 1**), the optimized toggle switch controls the expression of the *arsR H48A* chelator gene, enabling the bacteria to activate arsenite capture in response to an arsenite exposure.

Before assessing sequestration performance, we examined how this circuit affected cell physiology and growth. Using optical density measurements at 600 nm, we tracked the growth of engineered and wild-type *EcN* strains under both arsenite-free and arsenite-exposed conditions (**Figure 4A**). In the absence of arsenite, cells harboring the toggle switch circuit grew slightly slower than wild-type cells, suggesting that even the basal activity of the circuit imposed a small metabolic cost, which can be due to low-level chelator expression drawing on cellular resources. When exposed to arsenite, the growth rate of toggle switch-containing cells declined further, whereas control cells showed no significant change. This difference indicates that the reduced growth rate was not a direct toxic effect of arsenite, but rather a result of the circuit becoming activated and diverting energy toward chelator production.

**Figure 4.**
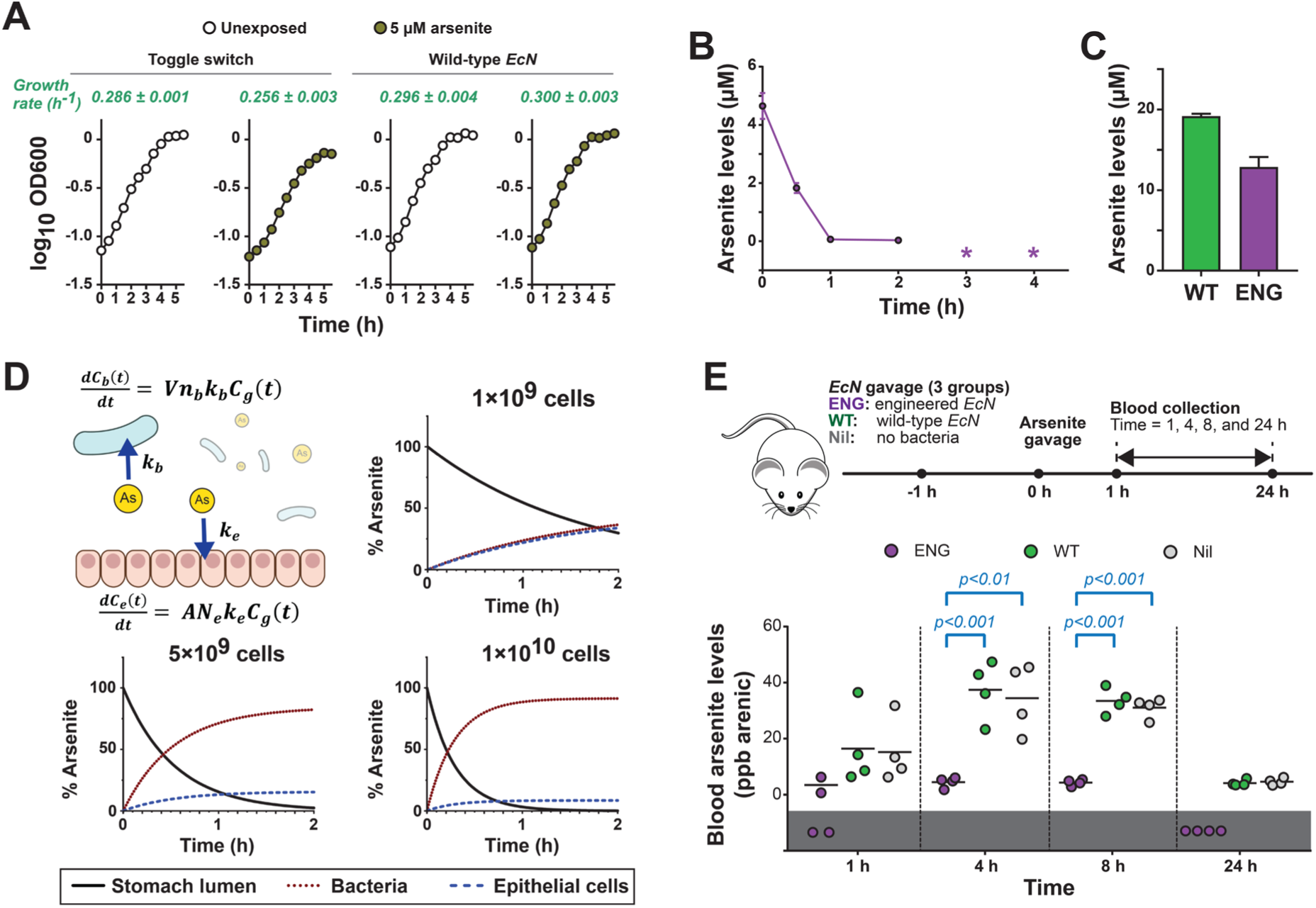
Characterization of engineered EcN for arsenite sequestration. (**A**) Growth of engineered *EcN* harboring the arsenite sense-and-response circuit (ENG) and wild-type *EcN* (WT) was measured by culture absorbance. Calculated growth rates for each strain in the presence of 0 or 5 µM sodium arsenite are shown above the corresponding plots. (**B**) In vitro arsenite sequestration by engineered *EcN* was quantified using ICP-MS. Data points represent the concentration of arsenite remaining in the culture supernatant after arsenite addition at time 0 h. An asterisk (*) indicates that arsenite levels were below the limit of detection at that timepoint. (**C**) Maximum sequestration capacity was assessed by exposing engineered and wild-type *EcN* cultures to 20 µM arsenite and analyzing supernatants after 3 h. For panels A to C, each data point represents the mean ± S.D. of three biological replicates; error bars are omitted when smaller than the marker. (**D**) A mathematical model was constructed to simulate arsenite mass transfer in the stomach lumen. Rate constants for arsenite uptake by engineered bacteria (*kb*) and epithelial cells (*ke*) were derived from experimental data and literature values. These constants are involved in defining the temporal contributions of bacterial uptake, *Cb(t)*, and epithelial uptake, *Ce(t)*, to changes in luminal arsenite concentration, *Cg(t)*. *Vnb* denotes the total number of bacterial cells, and *ANe* denotes the total number of epithelial cells. The model, detailed in the Methods, was used to simulate arsenite transfer during the first 2 h following oral gavage of engineered *EcN* and arsenite. (**E**) Blood arsenite levels in mice following bacterial treatment. The timeline (top) indicates the schedule of bacterial administration, arsenite exposure, and blood sampling. In the plot (bottom), each marker represents a data point from an individual mouse; bars denote mean values for four biological replicates. Markers within the gray shaded region indicate samples with undetectable arsenite. Two-way ANOVA revealed significant effects of engineered *EcN* treatment (F = 22; *p* < 0.001) and time (F = 56; *p* < 0.001). Significant pairwise differences identified by Tukey’s HSD are indicated in blue.

To further evaluate the physiological impact of circuit activation, we performed colony-forming unit (CFU) analyses (**Supplementary Figure S5**). After 3.5 hours of arsenite exposure, CFU counts of induced toggle switch cultures were roughly 10-fold lower than those of unexposed controls, consistent with a slowdown in cell growth. However, by 5.5 hours, CFU values reached above 10^9^ cells/mL, which is comparable to the other test conditions, demonstrating that the engineered *EcN* cells remained viable and capable of sustaining growth even after circuit activation, a critical property for its potential function within the gastrointestinal tract.

### In vitro characterization of arsenite sequestration

To reach our goal of using this engineered *EcN* to prevent arsenite absorption in a gastrointestinal tract, we first characterized its arsenite sequestration function in vitro, such that we can use these results to simulate the amount of bacterial cells required in the animal experiments for efficient sequestration. To determine the kinetics of arsenite sequestration in cultures (**Figure 4B**), we exposed cultures at the log phase (OD600 was 0.6 to 0.7) to 5 µM of sodium arsenite and collected samples to measure arsenite concentration in supernatant at different time points. After 0.5 hour of incubation, arsenite levels were reduced to 1.83 ± 0.17 µM; based on CFU analysis at the 0- and 1-hour time point, we estimated that there were about 4.6 ×10^9^ cells/mL in the culture for arsenite sequestration (**Supplementary Figure S6**). The initial sequestration rate was calculated as (1.22 ± 0.13) ×10^-18^ mol/cell/hour. The levels of arsenite decreased to below 0.1 µM after 1 hour and it was under our detection limit at the 3-hour time point.

Beyond sequestration kinetics, we also assessed the arsenite-binding capacity of the engineered *EcN* strain by challenging it with a higher arsenite concentration that exceeded its sequestration potential. Log-phase cultures were exposed to 20 µM sodium arsenite, and residual arsenite levels in the supernatant were measured at 0 and 3 hours. After 3 hours, arsenite concentrations decreased to 12.8 ± 1.0 µM (**Figure 4C**), corresponding to substantial uptake by the bacterial population. At this timepoint, cell density was estimated at (1.05 ± 0.41) ×10^10^ CFU/mL (**Supplementary Figure S6**), yielding an average arsenite sequestration capacity of (6.02 ± 0.97) ×10^-19^ mol/cell. These results quantify the per-cell arsenite retention ability of the engineered *EcN*, providing a useful benchmark for predicting the total detoxification potential achievable in vivo.

### Determination of the arsenite dose for mouse studies

After characterizing arsenite sequestration by the engineered bacteria in vitro, we next evaluated their ability to reduce arsenite absorption in a mouse gastrointestinal model. As a first step, we sought to identify an arsenite dose that would generate a detectable but unsaturated increase in blood arsenite levels. An ideal dose should be high enough to produce a measurable rise in blood concentration, yet low enough that a reduction in arsenite availability within the gut (such as through bacterial sequestration) would cause a proportional and significant decrease in blood arsenite levels. In contrast, at saturating doses, arsenite absorption becomes rate-limited, and changes in gut arsenite content would no longer be reflected in the bloodstream.

To determine such a dose, mice were orally gavaged with either 1 nmoles or 4 nmoles of sodium arsenite (**Supplementary Figure S7**). At the higher dose (4 nmoles), blood arsenite levels rose rapidly to 181 ± 66 ppb after 1 hour and continued increasing to 306 ± 25 ppb at 8 hours, remaining elevated (198 ± 44 ppb) even after 24 hours, indicating systemic saturation and incomplete clearance. In contrast, mice receiving 1 nmole of arsenite exhibited a more dynamic and responsive profile: blood arsenite reached 15 ± 11 ppb at 1 hour, peaked at 34 ± 12 ppb by 4 hours (remaining 31 ± 4 ppb at 8 hours), and returned near baseline (4.6 ± 1.2 ppb) at 24 hours.

Together, these results indicate that 1 nmol of arsenite provides an optimal dose for evaluating in vivo sequestration, as fluctuations in gut arsenite availability are expected to sensitively mirrored by changes in blood arsenite concentration.

### Computational simulations of arsenite transfer in the gastrointestinal model

Based on the in vitro sequestration parameters and the arsenite dose established for mouse studies, we developed a mathematical model to simulate arsenite mass transfer in the gastrointestinal tract. Our goal was to estimate the number of engineered *EcN* cells required to effectively prevent arsenite absorption (**Figure 4D**). The model tracks the time-dependent concentrations of arsenite in the stomach lumen, *C_g_(t)*; and the contributions of the loss of arsenite in the stomach lumen by engineered bacterial cells, *C_b_(t)* and the epithelial cells lining the stomach, *C_e_(t)*. The system is represented as a short, closed-segment interval with no external inflow or outflow and assumes no efflux, redistribution, or metabolism of arsenite by either bacterial or epithelial cells during the analysis window.

A mass balance is applied to the lumen and coupled to first-order uptake terms describing sequestration by bacteria (a volumetric uptake sink) and uptake by epithelial cells (a wall-associated sink). This formulation captures lumenal depletion of arsenite alongside its corresponding accumulation in the two biological sinks. The resulting system of equations, described in the Methods section, provides explicit solutions for arsenite concentrations in each compartment as functions of bacterial load and time. The bacterial uptake rate constant was derived from our experimental measurements (**Figure 4B**), while parameters describing the epithelial compartment, including stomach volume, surface area, and epithelial cell number, were obtained from the literature (*50–53*).

Using these inputs, the model simulated how arsenite distributes between epithelial and bacterial cells following oral gavage of 1 nmol arsenite in the presence of varying bacterial loads (1, 5, or 10 ×10^9^ cells). The simulations predicted that 1 ×10^9^ cells would sequester only ∼36% of arsenite within 2 hours, whereas 5 × 10^9^ cells would capture ∼82% (**Figure 4D**). Increasing the bacterial dose further to 1 ×10^10^ cells produced only modest additional benefit (∼91%).

Based on these simulations, we selected 5 ×10^9^ engineered cells as the dose for subsequent mouse experiments, balancing sequestration efficiency with the need to avoid unnecessary physiological stress from excessive bacterial loads.

### Effect of engineered *EcN* on blood arsenite levels

Building on these findings, we next evaluated whether the engineered *EcN* could protect the host from dietary arsenite exposure in vivo (**Figure 4E**). Mice were orally gavaged with a dose of engineered *EcN*; as controls, two additional groups received either wild-type *EcN* or no bacteria. One hour later, all mice were gavaged with 1 nmol of sodium arsenite, and blood samples were collected at 1, 4, 8, and 24 hours. At every time point, blood arsenite levels in the wild-type *EcN* and no-bacteria groups were comparable, indicating that unmodified *EcN* lacks endogenous pathways capable of reducing arsenite absorption through the gastrointestinal tract.

In contrast, mice treated with engineered *EcN* showed a pronounced reduction in arsenite uptake. Four hours after arsenite exposure, blood arsenite levels in the engineered *EcN* group were 4.5 ± 1.6 ppb, significantly lower than those in mice given wild-type *EcN* (37.4 ± 9.1 ppb) or no bacteria (34.5 ± 10.7 ppb). This protective effect persisted at the 8-hour time point, with engineered *EcN*-treated mice maintaining low arsenite levels (4.3 ± 1.0 ppb) compared to wild-type *EcN* (33.5 ± 4.0 ppb) and no-bacteria controls (33.1 ± 3.1 ppb). By 24 hours, arsenite was undetectable in the blood of mice treated with engineered *EcN*, whereas low but measurable levels remained in the wild-type *EcN* (4.2 ± 1.0 ppb) and no-bacteria (4.7 ± 1.0 ppb) groups.

Together, these results demonstrate that the engineered *EcN* substantially reduces arsenite absorption from the gastrointestinal tract, supporting the conclusion that sequestration occurred in situ and effectively prevented arsenite from entering the bloodstream.

## Discussion

### Important features in our strategy

In this study, we present a microbial sense-and-response strategy that enables selective sequestration of arsenite in the gastrointestinal tract before systemic absorption occurs. By engineering a gut bacterial strain with a synthetic toggle switch controlling expression of a protein-based arsenite chelator, we demonstrate an approach that is both preventative and targeted for managing dietary arsenic exposure. Our results in a mouse model show that a single dose of engineered bacteria before an arsenite exposure can markedly reduce entry of this heavy metal species into the bloodstream, establishing a functional proof of concept for probiotic-based detoxification.

A key innovation of our design is the integration of arsenite sensing with dynamically regulated chelator expression. Unlike conventional chelator therapies, which may remove both toxic and essential metal ions, our engineered bacteria activate chelator production only upon detection of arsenite. This conditional expression has several advantages. First, it minimizes off-target depletion of essential micronutrients, a major limitation of prophylactic chelation therapy (*54*). Second, it reduces the baseline metabolic burden on the bacteria, improving their fitness and persistence in the gastrointestinal tract. Third, the use of a toggle switch transforms a transient environmental signal into a sustained protective response, allowing cells to continue detoxifying even after arsenite levels decline—effectively extending the intervention window during the digestive process.

The H48A mutant of ArsR, developed here as a DNA-binding-deficient yet high-affinity arsenite chelator, further contributes to the efficiency and safety of this engineered activity. This design prevents unintended transcriptional repression of endogenous genes of the bacteria while retaining strong metal-binding capability. The resulting genetic modularity highlights a broader principle: protein-based chelation can be rationally tuned for performance, opening the door to engineering microbial systems that target other toxic metals or environmental contaminants.

Another unique feature of the system is its built-in growth-dependent shutoff mechanism. As the ON-state maintenance depends on cellular CelR levels that are diluted during active cell division, the circuit automatically resets under growth conditions. This behavior prevents long-term, unchecked chelator production in bacteria that may transiently colonize the gut—an important biosafety advantage for living therapeutics intended for repeated administration.

Finally, our combined experimental and mathematical modeling framework allowed us to determine bacterial dosing parameters needed for effective in vivo function. This quantitative approach provides a foundation for rational design of microbial detoxifiers and may be applicable to other xenobiotic sequestration problems.

With these features, our platform suggests a path toward practical, consumer-friendly strategies for mitigating chronic dietary exposure to toxic metals. As *EcN* is already used as a probiotic supplement in humans (*55*), an engineered version could, in principle, be formulated similarly, as capsules or suspensions taken daily or immediately before meals. Each dose would perform a single-pass detoxification by capturing heavy metals present in the ingested food and directing them toward fecal elimination, thereby reducing the cumulative body burden over time. This approach could be especially valuable in regions where groundwater and food sources are chronically contaminated with arsenite, or in populations for whom dietary control is difficult to maintain. Although translating such a product will require rigorous safety, stability, and regulatory assessments, the combination of selective activation, finite persistence, and defined dosing makes a daily-use formulation both conceptually feasible and potentially transformative as a preventative tool for environmental health.

### Limitations and challenges

While our results are encouraging, there are gaps and challenges that require to be overcome for robust applications. In our in vivo experiments, arsenite was administered only one hour after delivery of the engineered bacteria, when the bacterial concentration in the stomach is expected to remain high and the genetic circuit fully active. Under these conditions, we could robustly demonstrate proof-of-concept detoxification. However, further studies are required to evaluate how rapidly the engineered strain is cleared from the stomach, how long the cells remain in a physiologically active state, and how the protective effect decays over time. Future studies will need to quantify the temporal window of efficacy by tracking bacterial persistence, circuit activation dynamics, and sequestration capacity over extended periods following administration. Such data will be essential for informing dosing strategies, predicting real-world performance, and optimizing formulations aimed at routine or pre-meal use.

Additionally, regulatory concerns surrounding release, colonization, and clearance of genetically modified organisms must be considered (*56*). Although our growth-dependent shutoff offers an inherent safety mechanism, additional biocontainment systems or exclusively non-colonizing strains may be required for clinical deployment (*57, 58*).

## Materials and Methods

### Bacterial strains

For cloning and construction of genetic circuits, *E. coli XL1-Blue* was used as the plasmid host and cultured in Luria-Bertani (LB) medium. Screening and characterization of arsenite chelator genes were performed in *E. coli BL21-AI*, while characterization of genetic circuits was conducted in *E. coli Nissle* 1917 grown in M9 medium, which consisted of 1× M9 salts, 2 mM MgSO_4_, 0.1 mM CaCl_2_, 0.2% (w/v) glucose, and 0.2% (w/v) casamino acids. Unless otherwise specified, all cultures were incubated at 37 °C with shaking at 200 rpm in the presence of kanamycin (50 µg/mL). All chemicals and bacterial culture reagents were purchased from VWR (Radnor, PA) and Thermo Fisher Scientific (Waltham, MA).

### Mouse model

All animal procedures were approved by the University of North Texas Institutional Animal Care and Use Committee (Protocol #24007) and conducted in the Animal Vivarium at the University of North Texas.

Experiments were carried out using male C57BL/6 mice aged 7–8 weeks (Charles River, Wilmington, MA). Animals were housed in a specific pathogen–free barrier facility maintained at 30–70% humidity and 20–26°C, under a 12:12 h light–dark cycle. Mice were provided feed (Purina Conventional Mouse Diet, JL Rat/Mouse 6F Auto, #5K67) and water ad libitum. Each cage contained four mice, which served as four biological replicates for an experimental condition.

### Plasmid construction

Cloning was performed using standard molecular biology techniques with reagents from New England Biolabs (Ipswich, MA), including Phusion™ High-Fidelity DNA Polymerase, T4 DNA ligase, and the restriction enzymes listed below. The sequence of plasmid selected in each step was deposited in GenBank database.

To construct the control circuit in which ArsR regulates mCherry expression (**Figure 2A**), we used plasmid *pTR* from our previous study (*59*) as the backbone. This plasmid contains a kanamycin resistance cassette and a *ColE1* origin of replication. A synthetic DNA fragment containing the *arsR* and mCherry genes, along with their respective promoters and terminators, was inserted into *pTR* via the *PstI* and *MluI* restriction sites using Golden Gate assembly with the endonuclease BsaI-HFv2. These two restriction sites were eliminated at the ligation junction. The sequence of the resulting plasmid has been deposited in GenBank.

The arsenite-responsive toggle switch (**Figure 2A** and **Supplementary Figure S1**) was constructed from this control circuit by inserting a fragment containing the *celR* gene driven by the *P_LarsO_* promoter and its downstream promoter through the *SalI* and *AflII* sites. Expression of CelR was subsequently optimized by varying its ribosome binding site (RBS) between the *NheI* and *SacI* restriction sites. The optimized toggle-switch circuit sequence is available in GenBank.

To enhance output expression, we next incorporated T7 RNA polymerase into the toggle switch (**Figure 2E** and **Supplementary Figure S2**). The T7 RNA polymerase gene fragment replaced mCherry via the *AvrII* and *XhoI* restriction sites. A T7 promoter-mCherry-terminator fragment was then introduced using Golden Gate assembly with endonuclease BsaI-HFv2 to process the insert, which was ligated into the plasmid via *SalI* and *NcoI* (this *NcoI* site was not present in the product upon ligation). This mCherry cassette served as the reporter to characterize T7 polymerase activity. We further optimized T7 RNA polymerase expression by swapping its RBS between the KasI and AvrII sites. The finalized plasmid sequence has been deposited in GenBank.

This optimized circuit was subsequently used to drive chelator expression by replacing *mcherry* with the *arsR H48A* gene fragment via *NcoI* and *HindIII*, yielding the final construct shown in **Figure 1** (Accession Number PX644048).

To construct the arabinose-inducible system for expressing chelator derivatives (**Figures 3C** and **1D**), the plasmid *pET28b(+)* (*60*) was modified to generate a new vector, designated *pETnoL*. The original *pET28b(+)* plasmid contains a *T7* promoter with a *lac* operator for transcriptional regulation by LacI. To eliminate LacI-mediated repression, the lac operator was removed by replacing the promoter region between the *BglII* and *XbaI* restriction sites, resulting in a *T7* promoter unresponsive to LacI. The *arsR* gene was subsequently cloned into *pETnoL* via *NcoI* and *HindIII* restriction sites. The resulting plasmid, *pETnoL-arsR* served as the template for generating ArsR variant constructs through site-directed mutagenesis using the QuikChange™ method.

### Screening of Toggle Switch circuit variants

To identify the toggle switch variant with the desired response behavior (**Supplementary Figure S1**), each *EcN* strain harboring a circuit variant was grown overnight in M9 medium and then diluted 100-fold into 3 mL of fresh medium containing either 0 or 5 µM sodium arsenite. Cultures were incubated for 24 hours, after which samples were collected for flow cytometry analysis to quantify cellular mCherry fluorescence.

Cultures exposed to 5 µM arsenite were then centrifuged to collect the cell pellet, washed once, and resuspended in 3 mL of fresh medium without arsenite, followed by an additional 24-hour incubation before a second mCherry measurement was performed.

### Characterization of Toggle Switch

To assess circuit sensitivity to arsenite (**Figure 2B**), overnight cultures of engineered *E. coli* were diluted 100-fold into 3 mL of fresh M9 medium containing a range of sodium arsenite concentrations (0, 0.16, 0.31, 0.63, 1.25, 2.5, or 5 µM). Cultures were incubated under standard growth conditions for 24 hours, after which cellular mCherry fluorescence was quantified by flow cytometry. Data were analyzed using GraphPad Prism v7.05 and fitted to a nonlinear regression dose-response curve described by the equation:

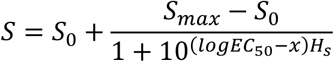

where *S_0_* and *S_max_* represent the basal and maximal mCherry fluorescence, respectively; S is the observed fluorescence signal; x is the logarithm of arsenite concentration; and EC_50_ and H_s_ (Hill slope) are constants derived from the fit.

To characterize the kinetics of output expression in response to arsenite (**Figure 2C**), mid-log phase cultures (OD_600_ ≈ 0.6) were exposed to 5 µM sodium arsenite. Samples were collected at 0, 0.5, 1, 2, 3, and 4 hours post-induction, and cellular mCherry fluorescence was quantified by flow cytometry.

The specificity of signal detection was assessed by diluting overnight cultures of *EcN* harboring the toggle switch plasmid 100-fold into fresh M9 medium supplemented with one of 16 test chemicals, as shown in **Figure 2D**. Cultures were incubated under standard growth conditions for 24 hours before flow cytometric analysis to determine cellular mCherry fluorescence.

To evaluate the stability of output expression under biostatic conditions (**Figure 2E**), overnight cultures of cells carrying the toggle switch circuit were first diluted 100-fold and then induced with 5 µM sodium arsenite. After overnight incubation (3 mL cultures), cellular mCherry levels were measured with flow cytometry; each culture was centrifuged to collect a cell pellet, which was washed once, and resuspended in an equal volume of fresh M9 medium to continue the incubation. This flow cytometric analysis and medium exchange were repeated every 24 hours for 3 days (72 hours total).

For experiments assessing the effect of cell growth on output expression (**Supplementary Figure S2**), induced overnight cultures were also washed and resuspended in fresh M9 medium as described above (**Figure 2E**); additionally, cultures were diluted 2-, 3-, or 6-fold after each resuspension by mixing 1.5, 1, or 0.5 mL of the culture, respectively, with fresh medium to a final volume of 3 mL. Samples were collected before each medium exchange (0, 24, 48, and 72 hours after arsenite removal) for flow cytometric analysis.

To characterize toggle switch performance when coupled to the T7 expression system (**Supplementary Figure S3** and **Figure 2F**), mid-log phase cultures were exposed to 0 or 5 µM sodium arsenite, and cell samples were collected after 1 and 24 hours of induction for fluorescence measurement by flow cytometry.

### Cellular fluorescence measurement by flow cytometry

To prepare samples for flow cytometry, 1 to 2 µL of cell culture was diluted in 150 µL of phosphate-buffered saline (PBS). The samples were analyzed using an ACEA NovoCyte 3000VYB flow cytometer (ACEA Biosciences, Inc.). For each experiment, at least 10,000 events were collected and gated based on forward and side scatter to exclude cell debris and aggregates. The geometric mean of fluorescence intensity for each sample was calculated using NovoExpress software (version 1.4.1; ACEA Biosciences, Inc.).

### Analysis of ArsR/SmtB family regulators

The NolR-DNA structure in **Figure 3A** was studied with the software PyMol v2.5. Sequence alignment (**Supplementary Figure S4**) was performed with an online tool, Clustal Omega (https://www.ebi.ac.uk/jdispatcher/msa/clustalo).

### Engineering of arsenite chelator

To evaluate the cytotoxicity of ArsR derivatives (**Figure 3B**), each *pETnoL* plasmid harboring an *arsR* mutant gene was transformed into *E. coli BL21-AI*, which carries an arabinose-inducible T7 RNA polymerase gene in its genome. Thus, arabinose addition induces ArsR expression in these cells. Cells containing a *pETnoL* plasmid without arsR gene were used as the control. Overnight cultures in LB medium were diluted 1:100 into 3 mL of fresh medium and grown to an OD_600_ of 0.5. A 10 µL aliquot from each culture was collected for colony-forming unit (CFU) analysis prior to induction, after which arabinose was added to a final concentration of 0.2% (w/v) to induce expression. Following 3 hours of induction, another 10 µL sample was taken for CFU determination. For CFU analysis, each sample was serially diluted 10-fold in phosphate-buffered saline (PBS), and 10 µL of each dilution was spotted onto LB agar plates containing kanamycin. Plates were incubated at 37 °C to allow colony formation.

To assess arsenite sequestration (**Figure 3C**), each strain was cultured in 5 mL of LB medium until reaching an OD_600_ of 0.5. Each culture was then divided into two 2-mL aliquots; one supplemented with 0.2% (w/v) arabinose (induced) and the other without arabinose (uninduced). After 3 hours of incubation, cells were harvested by centrifugation, washed, and resuspended in PBS containing 20 µM sodium arsenite. Following a 1-hour incubation, supernatants were collected for arsenite quantification by atomic absorption spectroscopy (AAS).

### Arsenite quantification by atomic absorption spectroscopy

Arsenite concentrations were quantified using a Thermo Fisher Scientific iCE™ 3500 Atomic Absorption Spectrometer equipped with a GFS35Z graphite furnace/autosampler. Data acquisition, standard curve fitting, and quantification analysis were performed using SOLAAR v3.0 software. Samples (**Figure 1D**) were diluted 8-fold with fresh PBS, and 1 mL of the diluted sample was mixed with 10 µL of TraceMetal™ grade nitric acid (Thermo Fisher Scientific; Cat. No. A509P500). For calibration, two stock solutions (0 and 5 µM sodium arsenite) were prepared in 1:100 nitric acid/PBS (v/v) and used by the instrument to generate standard concentrations of 0, 0.075, 0.15, 0.325, 0.625, 1.25, 2.5, and 5 µM arsenite; results from these standards were used to plot a quadratic standard curve for determine arsenite concentrations in biological samples. During data acquisition, the instrument automatically mixed 20 µL of each sample with 5 µL of the matrix modifier Specpure™ nickel nitrate (Thermo Fisher Scientific; Cat. No. AA39043AE) and processed them according to the graphite furnace parameters listed in **Table 1**. Absorbance was measured for 3 seconds in the Atomization phase at 193.7 nm with a band-pass of 0.5 nm, lamp current set to 90%, and Zeeman background correction applied.

**Table 1:** Graphite furnace parameters for arsenite quantification.

| Phase | Temperature (°C) | Hold time (sec) | Ramp (°C/sec) | Gas type | Gas flow (L/min) |
| --- | --- | --- | --- | --- | --- |
| Drying | 100 | 30 | 10 | Argon | 0.2 |
| Ashing | 1300 | 20 | 150 | Argon | 0.2 |
| Atomization | 2600 | 3 | Max. rate | N/A | Off |
| Tube clean | 2700 | 3 | Max. rate | Argon | 0.2 |

### Characterization of cell growth

To evaluate how the arsenite-sequestration circuit affects cellular fitness, we compared the growth of engineered *EcN* containing the toggle switch-arsenite chelator circuit to that of a wild-type *EcN* strain without the plasmid (**Figure 4A**); the wild-type strain did not have the kanamycin-resistance marker and it was grown in M9 media without the antibiotic. Overnight cultures were diluted 1:100 into 5 mL of fresh M9 medium and allowed to grow for 1 hour under standard conditions. Sodium arsenite was then added to a final concentration of either 0 or 5 µM, marking the 0-hour time point. At 0, 0.5, 1, 1.5, 2, 2.5, 3, 3.5, 4, 4.5, 5, and 5.5 hours post-addition, 200 µL of culture was transferred into a 96-well plate, and absorbance at 600 nm was measured using a Synergy H1 microplate reader (Agilent Technologies). Growth rates were calculated using the following equation:

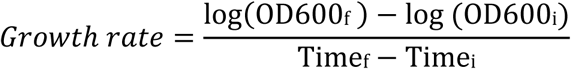

where OD600_f_ and Time_f_ correspond to measurements at the 4-hour time point, and OD600_i_ and Time_i_ correspond to the 0.5-hour time point. These time points were selected because the cultures exhibited clear exponential growth during this interval. To assess viable cell density, samples were also collected at 0.5, 3, and 5.5 hours for CFU analysis (**Supplementary Figure S5**).

### In vitro characterization of arsenite sequestration

To quantify the rate of arsenite sequestration (**Figure 4B**), engineered *EcN* cultures were grown in 5 mL of M9 medium to an OD600 of approximately 0.7. Sodium arsenite was then added to a final concentration of 5 µM. At 0, 0.5, 1, 2, 3, and 4 hours post-addition, 300 µL of each culture was collected, centrifuged, and the supernatant was analyzed by inductively coupled plasma mass spectrometry (ICP-MS) to determine arsenite concentrations. CFU measurements were performed at 0, 1, 2, 3, and 4 hours to estimate viable cell density (**Supplementary Figure S6**).

To determine the arsenite sequestration capacity (**Figure 4C**), cultures at OD600 ≈ 0.7 were exposed to 20 µM sodium arsenite, and samples were collected at 0 and 3 hours for ICP-MS analysis. CFU counts were obtained at both time points (**Supplementary Figure S6**).

### Mass transfer simulations

We developed a mathematical model to simulate the transfer of arsenite from the stomach lumen into engineered bacteria and gastrointestinal epithelial cells (**Figure 4D**). To simplify the system, we assumed that: (i) neither arsenite nor bacterial cells exit the stomach compartment; (ii) arsenite internalized by cells does not return to the lumen; and (iii) the total amounts of arsenite, engineered bacteria, and epithelial cells remain constant over time. Under these assumptions, we described the time-dependent distribution of arsenite across the three compartments using the following equations:

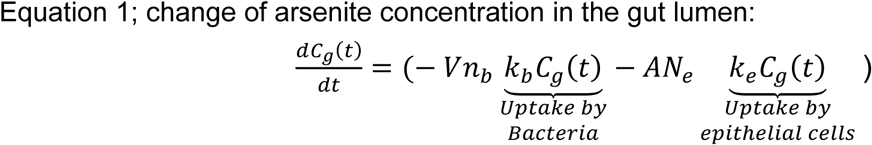

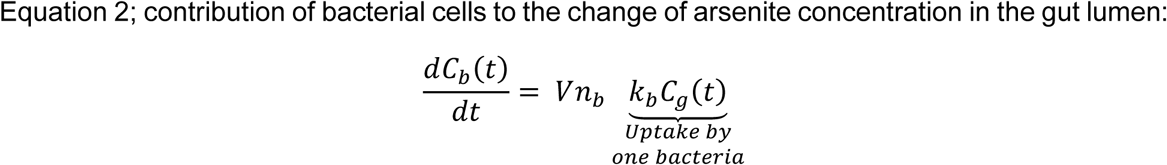

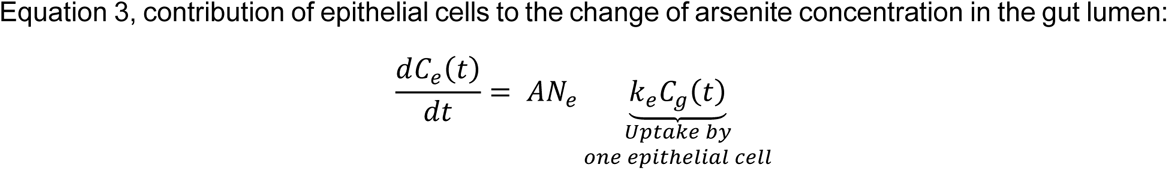

Where *C_g_(t)* is the concentration of arsenite (µM) in the stomach lumen, and *C_b_(t)* and *C_e_(t)* are the contributions of bacterial cells and epithelial cells, respectively, to the decrease in arsenite in the stomach lumen; *k_b_* and *k_e_* are bacteria and epithelial cell uptake rate coefficient; *V* is stomach volume, *n_b_* is bacterial cell density; *A* is the gut interior wall area; and *N_e_* is the number of epithelial cells per unit area of gut wall.

By solving these equations, we derive explicit expressions (Equations 4 to 6) that predict the temporal concentration of arsenite in lumen, bacteria, and epithelial cells as a function of bacterial cell number.

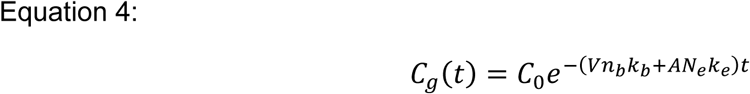

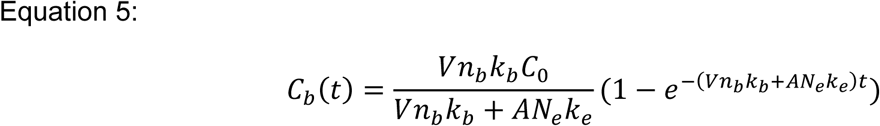

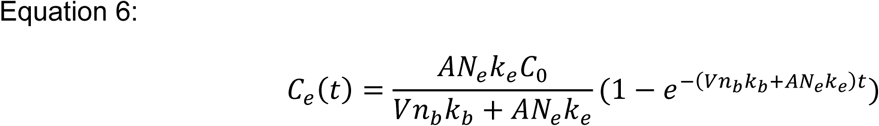

In these equations, the bacterial cell density, *n_b_* (cells mL^-1^), and time, *t* (hour), are variables.

The values of all epithelial cells-related parameters were obtained from the literature: gut surface area, *A*, is 1.4 ×10^6^ mm² (*61*). Lumen volume, V, is 0.4 mL (*52*). Arsenite uptake rate coefficient of epithelial cells is reported as 0.11 h^-1^ (*50*); in this study, it was normalized on a per-cell, per-hour basis to yield *k_e_* = 0.11 ×10^-10^ h^-1^ cell^-1^. The epithelial cell density on the gut wall, *N_e_*, was estimated to be 1.9 ×10^4^ cells mm^-2^ based on published data (*53*).

The arsenite uptake rate coefficient, *k_b_*, was estimated from our experimental data in **Figure 4B** with the following equation:

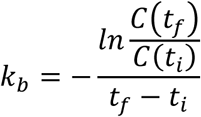

Where *C(t)* is the concentration of arsenite in the supernatant at time t. We used data from timepoints 1-hour and 2-hour to calculate *k_b_*, which resulted as 6.3 × 10^-10^ cell^-1^ h^-1^.

These parameters were then used to compute values of *C_g_(t)*, *C_b_(t)*, and *C_e_(t)* by using MATLAB R2023b. Results with bacterial cells number, *Vn_b_* = 1, 5, and 10 ×10^9^ cells were plotted (**Figure 4D**).

### Animal studies

All mice were quarantined in the animal facility for at least one week prior to experimentation. To characterize dose-dependent arsenite absorption (**Supplementary Figure S7**), each mouse received 100 µL of sodium arsenite solution (10 or 40 µM) by oral gavage, corresponding to a total dose of 1 or 4 nmol of arsenite. Blood samples (100 µL) were collected from the tail vein using a 25-gauge needle at 1, 4, 8, and 24 hours after gavage. To prevent clotting, each sample was immediately mixed with 10 µL of 2% (w/v) EDTA. Samples were then centrifuged to isolate plasma for ICP-MS analysis.

To assess the effect of engineered *EcN* on arsenite absorption (**Figure 4E**), *EcN* cultures were grown in M9 medium to an OD600 of ∼0.7, corresponding to an estimated density of ∼5 ×10^9^ cells/mL. For each mouse, 1 mL of culture was centrifuged, and the resulting pellet was resuspended in 100 µL of PBS for oral gavage. One hour later, mice received arsenite, and blood collection proceeded, following the same protocol used in the dose-dependent absorption study (**Supplementary Figure S7**).

Statistical analysis of animal studies was performed with the software, GraphPad Prism v7.0.5. A two-way ANOVA was conducted with treatment and time as independent variables and blood metal levels as the dependent variable, testing for the two main effects (treatment and time) and their interactions to determine whether the temporal trajectory of blood metal levels differed across groups. Significant effects were followed by post hoc pairwise comparisons with correction for multiple testing (e.g., Tukey’s HSD) to identify specific time points and treatment groups that differ.

### Inductively Coupled Plasma Mass Spectrometry

ICP-MS analysis was performed with an Agilent LC-linked ICP-MS 7850 with dual autosamplers (Agilent Technologies). It was operated under the following optimized conditions: radio frequency (RF) power of 1550 W, plasma gas flow of 15 mL/min, auxiliary gas flow of 0.9 mL/min, and nebulizer gas flow rate of 1.08 L/min. These parameters ensured stable plasma conditions, efficient ionization of analytes, and high analytical sensitivity. For sample preparation, 50 µL of each blood plasma sample was mixed with 5 mL of a diluent containing 0.5% HNO_3_ and 2% ethanol (v/v). The diluted mixture was vortexed thoroughly to ensure complete homogenization and then analyzed on the same day to minimize degradation. Matrix-matched calibration solutions were prepared from the multi-element stock standard (Inorganic Ventures, Christiansburg, VA) in 0.5% HNO_3_ and 2% ethanol (v/v), using pooled blood plasma from exposed mice to account for matrix effects of blood analysis with calibration levels set at 0.01, 0.1, 0.5, 1, 2.5, 5 and 15 µg/L. An internal standard solution containing 1 µg/mL yttrium (Y), was continuously introduced into the system throughout the analysis. The regression coefficient of the calibration curve above 0.995, was accepted for quantification. The detection limit for arsenic was 0.018 µg/L (2.34 ×10^-4^ µM).

### Graphical illustrations

We used two images from NIH BioArt Source to create figures, including [NIAID Visual & Medical Arts. (10/7/2024). Bacillus Bacteria. NIAID NIH BIOART Source. bioart.niaid.nih.gov/bioart/43]; and [NIAID Visual & Medical Arts. (10/7/2024). Basal Cell Brush. NIAID NIH BIOART Source. bioart.niaid.nih.gov/bioart/45].

## Supplementary Materials

One supplementary file is included in this submission:

Supplementary Materials**—Containing** Supplementary Figures S1 to S7

## Acknowledgements

We thank Dr. Brianne Soulen and the UNT Advanced Environmental Research Institute for guidance in performing atomic absorption spectroscopy analyses. We also acknowledge the use of the online tool ChatGPT 5.1 for assistance with grammar and text refinement.

## Author contributions

Conceptualization: CTYC

Methodology: NN, MW, SSM, VK, FE, LJS, LL, and CTYC

Investigation: NN, MW, SSM, VK, and FE

Visualization: FE and CTYC

Supervision: FE, LJS, LL, and CTYC

Writing—original draft: CTYC

Writing—review & editing:

## Data and Materials Availability

All data needed to evaluate the conclusions in the paper are present in the paper and/or the Supplementary Materials. Sequences of plasmids developed in this study are deposited in GenBank database:

*GenBank will release them to the public database until the accession numbers appear in print.*

