## Supplementary Figure for "Engineered probiotics that sequester arsenite in a mouse gastrointestinal system"

**Authors and Affiliations:**

Nhu Nguyen [1,‡], Miaomiao Wang [1,‡], S. Sifiso Msibi [2], Vincenzo Kennedy [1], Fateme Esmailie [1], L. Joseph Su [2], Lin Li [1], Clement T. Y. Chan [1,3,*]

[1] Department of Biomedical Engineering, University of North Texas, Denton, TX, USA

[2] Peter O'Donnell Jr. School of Public Health, UT Southwestern Medical Center, Dallas, TX, USA

[3] BioDiscovery Institute, University of North Texas, Denton, TX, USA

[‡] These authors are considered to have equal contributions to this study

* Corresponding author

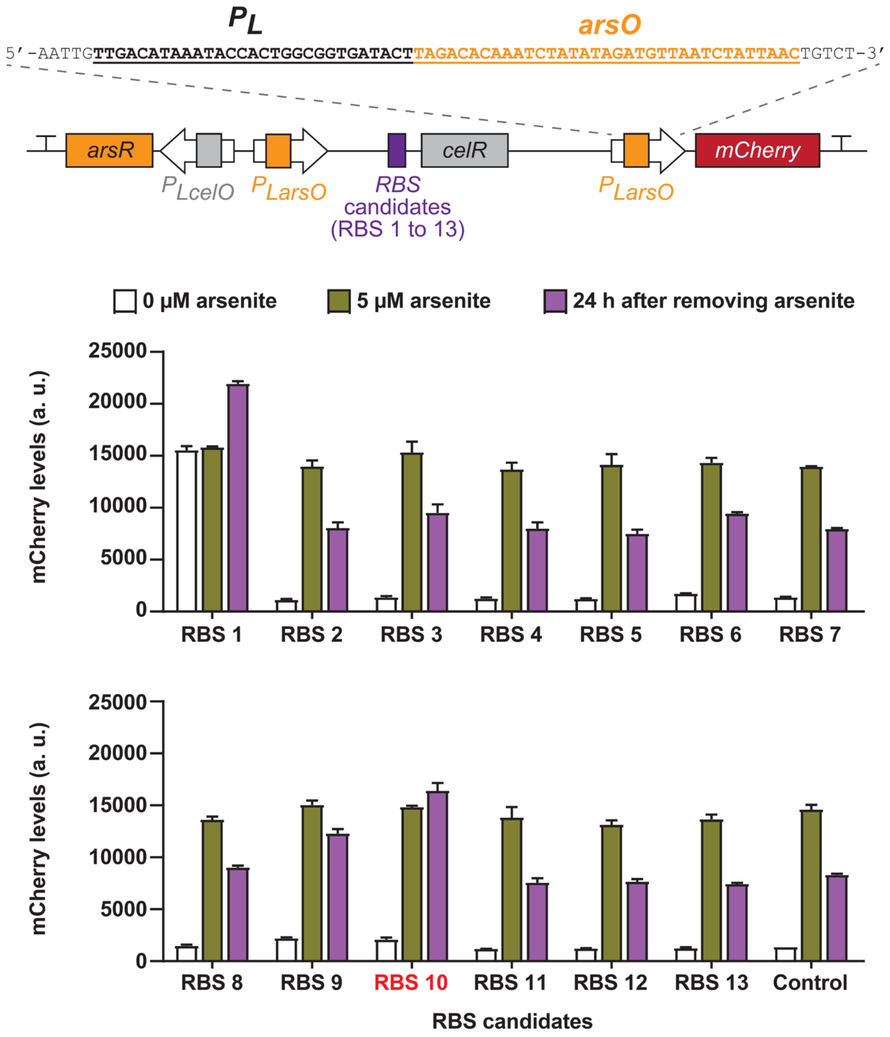


**Supplementary Figure S1. Screening of arsenite-responsive toggle switch.** In this circuit (top), a panel of ribosomal binding sites (RBSs) was screened for modulating *celR* expression to achieve desirable arsenite-responsive toggle switch behaviors. The ArsR-regulated promoter, $P_{LarsO}$, was constructed in this study, and its sequence is shown. *EcN* strains harboring individual circuit variants were grown overnight in the presence of either 0 or 5 μM sodium arsenite (indicated as "0 μM arsenite" and "5 μM arsenite," respectively), and cellular mCherry fluorescence was measured. To evaluate the ability of each variant to maintain output expression in the absence of arsenite during biostatic conditions, cultures initially exposed to 5 μM arsenite were washed and resuspended in fresh arsenite-free medium, then grown for an additional 24 h ("24 h after removing arsenite"); mCherry levels were then quantified. Each data point represents the mean ± S.D. of three biological replicates. The circuit variant with RBS 10 (colored red) was selected for continuing this study.

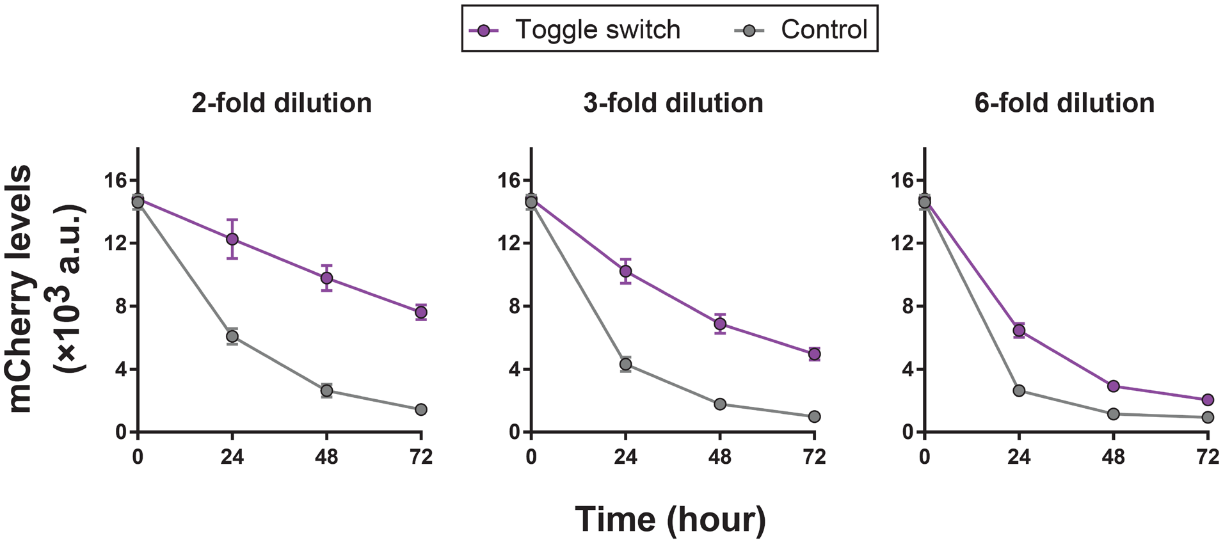


**Supplementary Figure S2. Characterization of output loss from the toggle switch during cell growth.** *EcN* cultures harboring the selected toggle switch variant (see Supplementary Figure S1) were induced overnight with 5 μM arsenite. Following induction, cells were washed, resuspended in fresh medium, and diluted 2-fold (left), 3-fold (middle), or 6-fold (right). Cellular mCherry fluorescence was measured after 24 hours, and this dilution-and-growth cycle was repeated until the 72-hour timepoint. Each data point represents the mean ± S.D. of three biological replicates.

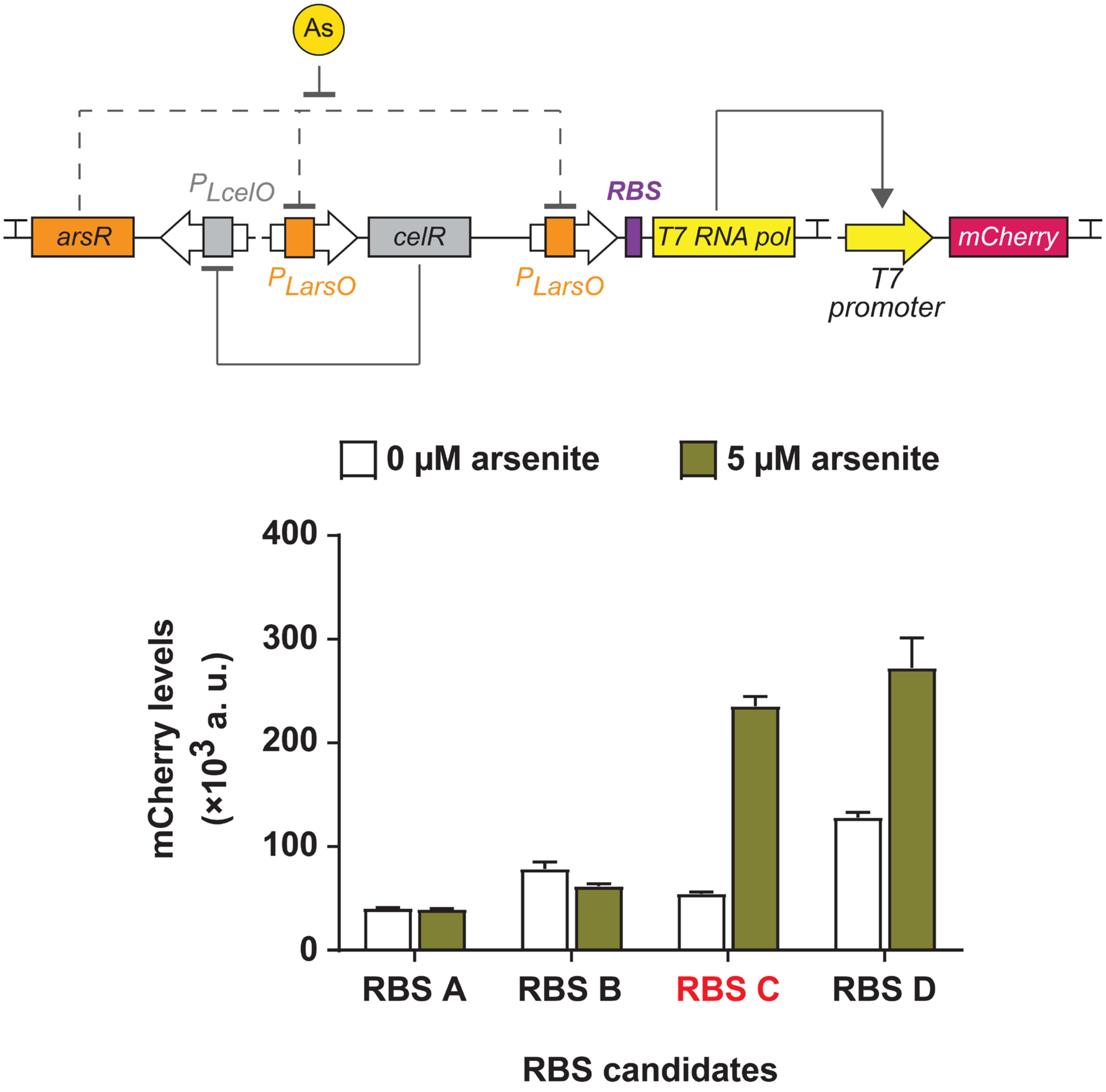


**Supplementary Figure S3. Optimization of T7 RNA polymerase activity to enhance output production.** In this circuit design, the toggle switch drives expression of T7 RNA polymerase to amplify downstream output expression. A series of ribosome binding sites (RBSs) was tested to tune T7 RNA polymerase activity. Mid-log phase *EcN* cultures carrying each circuit variant were exposed to 0 or 5 μM sodium arsenite for 1 hour, after which cellular mCherry fluorescence was measured. Each data point represents the mean ± S.D. of three biological replicates. The variant incorporating RBS C to control T7 RNA polymerase expression was selected for subsequent studies.

```
RzcA (O85142)            ---MSEQYSEINTDTLERVTEIFKALGDYNRIRIMELLSVSEASVGHISHQLNLSQSNVS 57
NolR (Q83TD2)            MEHAMQPLSPEKHEEAEIAAGFLSAMANPKRLLILDSLVKEEMAVGALANKVGLSQSALS 60
ArsR (Q01256)            -------------MSYKELSTILKVLSDPSRLEILDLLSCGELCACDLLEHFQFSQPTLS 47
                                         :  : ::..:.: .*: *:: *   * ..  : .:. :**  :*

RzcA (O85142)            HQLKLLKSVHLVKAKRQGQSMIYSLDDIHVATMLKQAIHHANHPKESGL----------  106
NolR (Q83TD2)            QHLSKLRAQNLVSTRRDAQTIYYSSSSDSVMKILGA-LSEIYGAATSVVIEKPFVRKSA  118
ArsR (Q01256)            HHMKSLVDNELVTTRKNGNKHMYQLNHEFLDYINQN-LDIINTSDQGCAC-KNMKSGEC  104
                         :::. *   .**.::::.:.  *. .   :  :    :        .
```

**Supplementary Figure S4. Sequence alignment of AsrR/SmtB family regulators.** Three family members were analyzed; their gene name and Uniprot ID were shown. The region of AsrR mutated is highlighted in purple, which is homologous to the α4 region of NolR that interacts with the DNA major groove as shown in Figure 3A.

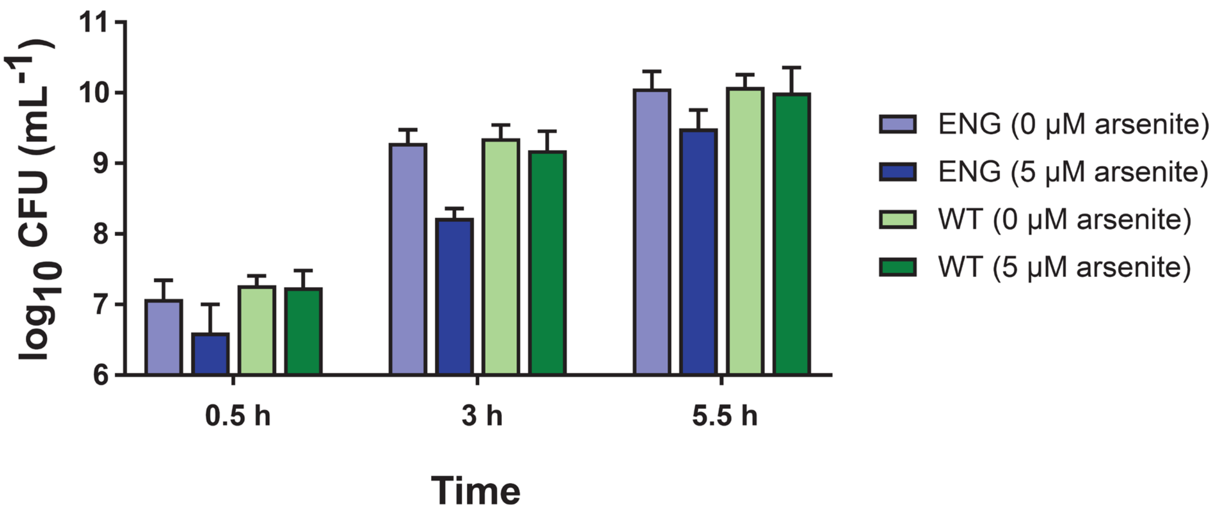


**Supplementary Figure S5. Colony forming unit (CFU) analysis of engineered *EcN* with the arsenite sense-and-response circuit.** Cultures of engineered *EcN* containing the toggle switch-arsenite chelator circuit (ENG) and the wild-type *EcN* (WT) were exposed to 0 and 5 µM sodium arsenite at OD600 ~ 0.1 (experiments described in Figure 4A). Viable cell density was quantified by CFU analysis at 0.5, 3, and 5.5 hours after exposure. Each data point represents mean ± S.D. of three biological replicates.

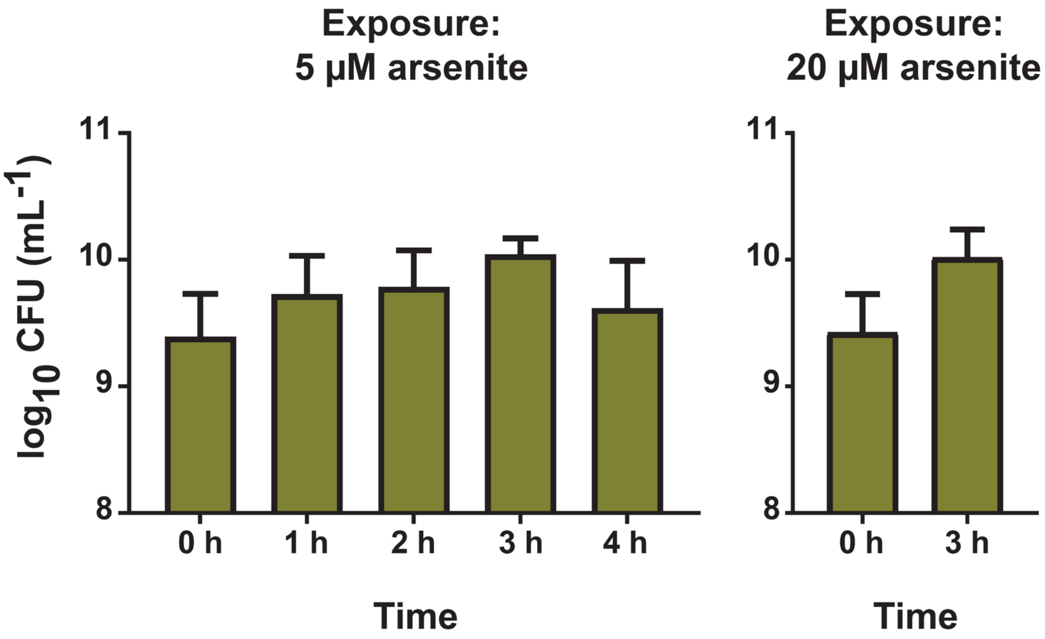


**Supplementary Figure S6. Cell viability of engineered *EcN* during in vitro arsenite sequestration.** Colony-forming units (CFUs) were quantified to assess cell viability during the sequestration experiments shown in Figure 4. The left plot reports CFUs from cultures exposed to 5 µM arsenite over a 0 to 4 h period (corresponding to Figure 4B). The right plot shows CFUs from cultures treated with 20 µM arsenite for 0 and 3 h (corresponding to **Figure 4C**). Each data point represents the mean ± S.D. of three biological replicates.

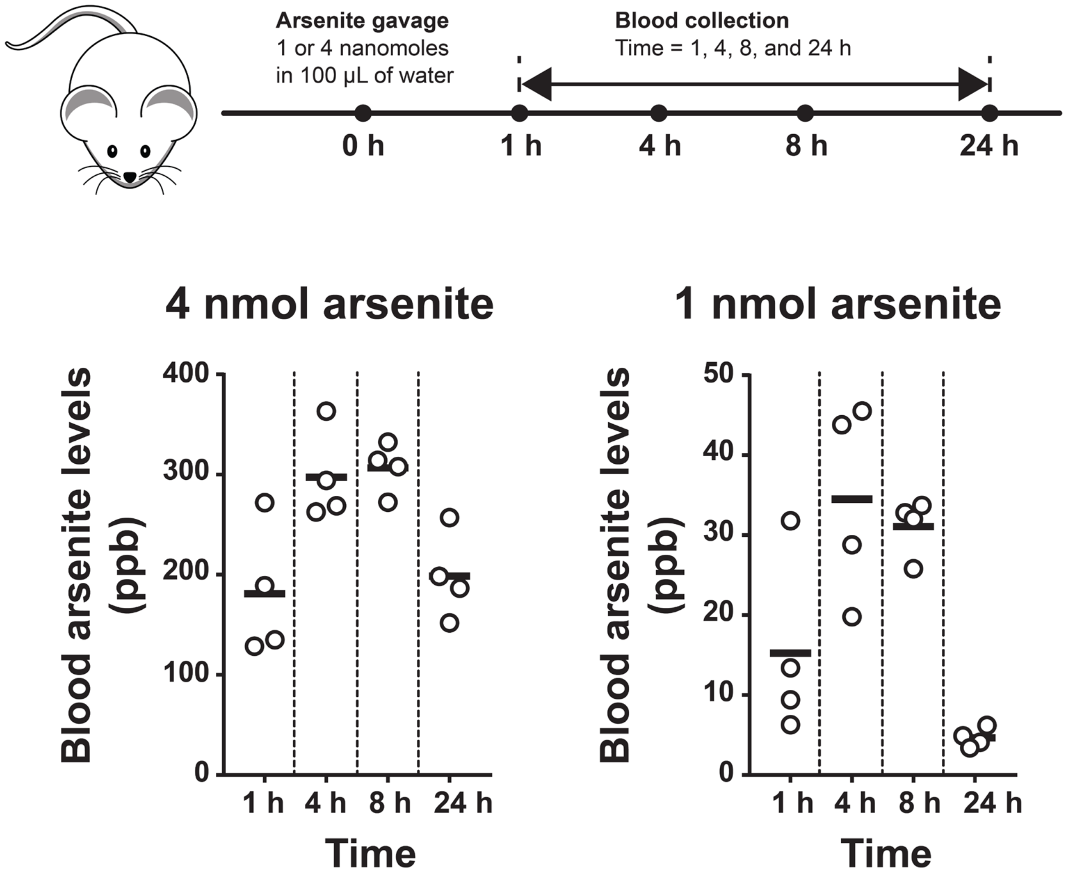


**Supplementary Figure S7. Blood arsenite levels in a mouse model following high- and low-dose oral gavage.** Mice were orally administered either 4 or 1 nmol of sodium arsenite, and blood samples were collected at 1, 4, 8, and 24 hours to quantify circulating arsenite levels. Each data point represents the measurement from an individual sample, and each bar indicates the mean value from four mice per group.
